# Horizontal Transfers and Gene Losses in the phospholipid pathway of *Bartonella* reveal clues about early ecological niches

**DOI:** 10.1101/003350

**Authors:** Qiyun Zhu, Michael Kosoy, Kevin J. Olival, Katharina Dittmar

## Abstract

Bartonellae are mammalian pathogens vectored by blood-feeding arthropods. Although of increasing medical importance, little is known about their ecological past, and host associations are underexplored. Previous studies suggest an influence of horizontal gene transfers in ecological niche colonization by acquisition of host pathogenicity genes. We here expand these analyses to metabolic pathways of 28 *Bartonella* genomes, and experimentally explore the distribution of bartonellae in 21 species of blood-feeding arthropods. Across genomes, repeated gene losses and horizontal gains in the phospholipid pathway were found. The evolutionary timing of these patterns suggests functional consequences likely leading to an early intracellular lifestyle for stem bartonellae. Comparative phylogenomic analyses discover three independent lineage-specific reacquisitions of a core metabolic gene - NAD(P)H-dependent glycerol-3-phosphate dehydrogenase (*gpsA*) - from Gammaproteobacteria and Epsilonproteobacteria. Transferred genes are significantly closely related to invertebrate *Arsenophonus*-, and *Serratia*-like endosymbionts, and mammalian *Helicobacter*-like pathogens, supporting a cellular association with arthropods and mammals at the base of extant bartonellae. Our studies suggest that the horizontal re-aquisitions had a key impact on bartonellae lineage specific ecological and functional evolution.

## Introduction

The increased availability of whole-genome data is providing more comprehensive insights into microbial evolution (Toft and Andersson 2010). One phenomenon of bacterial evolution concerns a process known as horizontal gene transfer (HGT), where bacteria transfer genetic material to related or to unrelated bacterial lineages (Doolittle 1999; Doolittle et al. 2003). From a biological perspective, HGT is vital for the origination of new bacterial functions, including virulence, pathogenicity or antibiotic resistance (Barlow 2009; Gophna et al. 2004; Koonin et al. 2001).

Instances of HGT also contain important information about evolutionary events in the bacterial lineage. Specifically, uneven distribution patterns of genes across lineages not only speak to the potential presence of HGT, but also to its frequency throughout lineage evolution. Depending on the evolutionary history post transfer, an HGT event may be informative about the directionality and mode of transfer, allowing identification of donor and recipient genomes.

Because many bacteria have niche preferences, the identification of the donating lineage may provide specific information about the nature of the environment the exchange took place in. This is particularly interesting for intracellular pathogens, as it implies HGT to have occurred in a specific environment - the host cells. The taxonomic identification of putative bacterial donors therefore allows inferences about the ancestral bacterial community composition at the time of exchange, although the extant host range and microbiome diversity may have changed.

With this in mind, we analyzed 28 currently available genomes of the bacteria *Bartonella* for HGT events. *Bartonella* species are gramnegative Alphaproteobacteria, and are thought to persist mainly as facultative intracellular invaders. They have been classified as emergent pathogens, and are ubiquitously associated to mammals, where they parasitize erythrocytes and endothelial cells (Pulliainen and Dehio 2012). More than half of the known *Bartonella* species are pathogenic to humans, and clinical manifestations vary from acute intraerythrozytic bacteremia to vasoproliferative tumor growth (Harms and Dehio 2012; Kaiser et al. 2011). Although it is known that bartonellae readily straddle the boundary between mammals and invertebrates, their ecological past remains obscure (Chomel et al. 2009). Phylogenetically, bartonellae form a derived monophyletic clade within the mostly plant associated Rhizobiales (Engel et al. 2011; Gupta and Mok 2007; Guy et al. 2013). Bartonellae have been increasingly detected in a broad range of blood-feeding or biting insects, and recent research on their diversity in blood-feeding insects suggests an early association to fleas (Morick et al. 2013; Tsai et al. 2011), but the full range of invertebrate associations is still underexplored.

Evidence for HGT in bartonellae has been found in previous studies, which mainly concentrated on the identification of gene transfer agents involved in the spread of known host adaptability and pathogenicity genes, including the T4SS secretion system (VirB, Trw, and Vbh) (Berglund et al. 2009; Saisongkorh et al. 2010). Surprisingly, little information exists on the horizontal transfer of other, more fundamental operational genes, which may also have implications for host-adaptation. Specifically, bartonellae are thought to be in the early stages of a transition to stable intracellularity (Toft and Andersson 2010). While they are stealthy pathogens in their mammalian hosts, and can survive and reproduce intra- and extracellularly, they have also been discussed as intracellular endosymbionts in their insect hosts (e.g. fleas, ked flies, bat flies) (Tsai et al. 2011). This transitional lifestyle has genomic ramifications, which have been associated with processes of gene loss, HGT, and recombination that specifically affect genes coding for cell membrane formation (outer surface structures), or intermediate metabolism (Toft and Andersson 2010; Zientz et al. 2004). Consequently, exploring the signature, provenance and order (timing) of HGT events in these pathways may be crucial in understanding the particular steps involved in the development of an intracellular lifestyle in bartonellae.

We here examine horizontal and vertical patterns in the evolution of a core metabolic pathway in bartonellae. Specifically, we employed comparative genomic analyses, phylogenetics and experimental approaches to explore the evolutionary successions of gains and losses of genes, with the goal to elucidate the ancestral and extant biological associations of bartonellae on organismal and cellular levels.

## Material and Methods

### Taxon sampling

Genomic data of *Bartonella* and other bacterial organisms related to this study were downloaded from the NCBI GenBank (http://www.ncbi.nlm.nih.gov/) or from the website of the Bartonella Group Sequencing Project, Broad Institute of Harvard and MIT (http://www.broadinstitute.org/). *Bartonella* species were grouped into four lineages (L1-L4) plus *B. tamiae* and *B. australis*, following the current taxonomy (Engel et al. 2011; Guy et al. 2013; Pulliainen and Dehio 2012). For the purpose of this paper we will refer to all bartonellae except *B. tamiae* as eubartonellae. This is based on the recognition that *B. tamiae* has been described as clearly distinct from all other currently known bartonellae lineages (Guy et al. 2013; Kosoy et al. 2008). A total of 28 *Bartonella* species were examined in this study (Table 1).

**Table 1.**
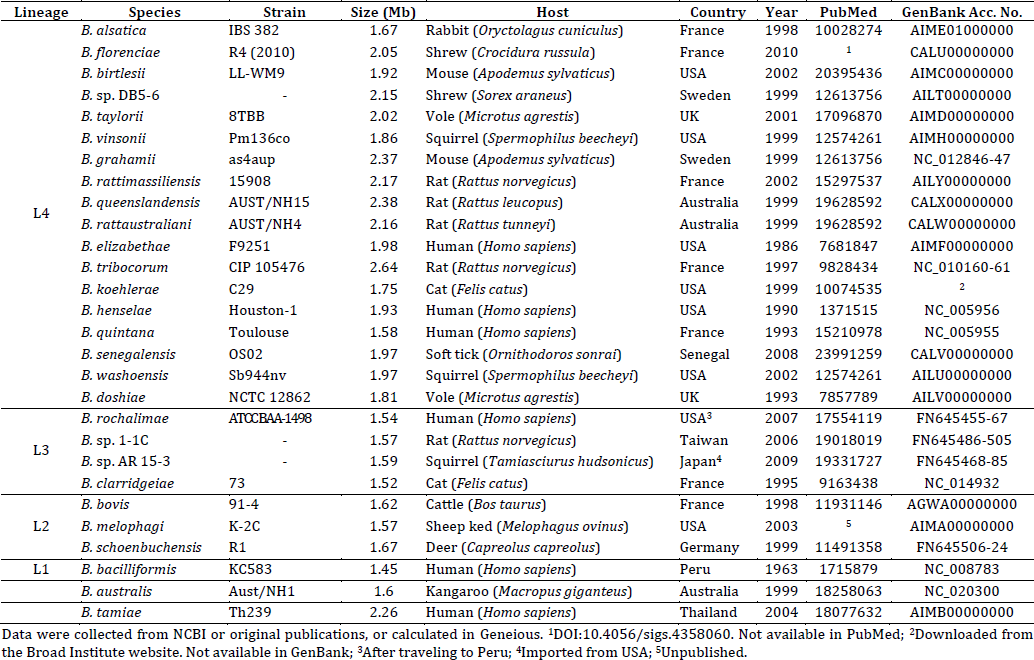
Basic information of *Bartonella* genomes and corresponding samples assessed in this study.

### BLAST hit distribution analysis of *Bartonella* genomes – initial discovery screen

Initial discovery analysis of putative HGT events in metabolic pathways was conducted under the assumption of aberrant BLAST hit distribution (Berglund et al. 2009; Koonin et al. 2001). Assisted by a homemade automated pipeline (available in the Dittmar Lab: https://github.com/DittmarLab/HGTector), batch BLASTP of *Bartonella* protein-coding genes was performed against the NCBI nr database (E-value cutoff = 1 × 10^−5^, other parameters remain default). Genes that have few or no hits from the close relatives of *Bartonella* (i.e., other Rhizobiales groups), but meanwhile have multiple top hits from taxonomically distant organisms (non-Rhizobiales groups), were considered to be candidates of HGT-derived genes and were subject to further phylogenetic analyses (see below). Particular attention was paid to genes involved in the core central intermediate metabolism, and cell wall formation, which have been identified in previous studies on bacterial metabolism (Zientz et al. 2004).

### Phylogenetic analyses of horizontally transferred genes

Phylogenetic analyses were employed to validate the putative HGT and vertical histories of the genes identified in the initial discovery screen. Phylogenetic patterns nesting a *Bartonella* gene within a homologous gene clade of a candidate donor group, or as strongly supported sister group of a candidate donor group, were considered significant evidence supporting the horizontal transfer from this particular donor to *Bartonella* (Husnik et al. 2013; Koonin et al. 2001; Nelson-Sathi et al. 2012; Schonknecht et al. 2013). Nucleotide sequences of genes of interest were extracted from *Bartonella* genomes as well as genomes of selected organisms that represent the putative donor group and its sister groups. Sequences were aligned in MAFFT version 7 (Katoh and Standley 2013), using the L-INS-i algorithm. The phylogenies of single gene families were reconstructed in a Bayesian MCMC statistical framework using MrBayes 3.2 (Ronquist et al. 2012), as well as a maximum likelihood (ML) method implemented in RAxML 7.7 (Stamatakis 2006). The Bayesian MCMC runs had a chain length of 20 million generations, with the sample frequency set as 1000. The optimal nucleotide substitution models for all three codon positions were computed in Partition-Finder 1.1 (Lanfear et al. 2012). Three independent runs were performed for each dataset to ensure consistency among runs. Trace files were analyzed in Tracer 1.5 (Drummond and Rambaut 2007) to check for convergence in order to determine a proper burn-in value for each analysis. A consensus tree was built from the retained tree-space, and posterior probabilities are reported per clade. The maximum likelihood (ML) was run implementing the GTR+G model (for all codon positions) and a bootstrap analysis was performed to gauge clade support.

### Survey of genomic environments

In order to determine the frequency, components and boundaries of the putatively horizontally transferred genetic material, genomic environments were manually examined in Geneious 6.1 (Biomatters). Our assumptions are that multiple independent transfers of a gene would likely result in different gene environments being affected. Likewise, if different *Bartonella* species share the same gene environment adjacent to horizontally transferred genetic material, and the transferred genes follow the previously detected vertical evolutionary pattern for bartonellae, presumably a single ancestral HGT event can be inferred for all species in that lineage. Putatively HGT-derived genes and their adjacent genomic elements were identified in recipient and donor genomes and compared across species and within lineages. Results from this analysis were mapped onto the *Bartonella* species tree (see below).

### Molecular evolution analyses

Selection analyses were carried out to gage selective pressures operating on all genes in the phospholipid pathway. Selection was assessed using the maximum likelihood (ML) method in the Codeml program of the PAML 4.7 package (Yang 2007). As the first step, an analysis under the one-ratio model (M0) was performed to estimate a global ω value (dN/dS ratio) across the phylogenetic tree. Global selective pressures were assessed using the site models (M1a, M2a, M8 and M8a). Evolutionary rates of particular branches of interest (ω_1_) versus the background ratio (ω_0_) were computed using the branch model (model=2). Selective pressures operating on subsets of sites of these branches were calculated using the branch-site models (model A and A1). The significance of change of ω value and evidence of positive selection was assessed using the likelihood ratio test (LRT). Positive sites were identified using the Bayes Empirical Bayes (BEB) analysis (Yang et al. 2005).

The tertiary structure of the GpsA protein in *Coxiella burnetii* (Gammaproteobacteria: Legionellales) was used to model the position of the identified sites with positive selection in horizontally transferred *gpsA* genes (Minasov et al. 2009; Seshadri et al. 2003).

The possibility that the horizontally acquired *gpsA* genes underwent convergent evolution in *Bartonella*, relative to their ancestors was explored. Potential ancestral states of the *gpsA* genes before HGT were reconstructed under Model A in Codeml of PAML (see above). The sequence was then compared to the consensus sequence of extant *Bartonella* species. A statistical approach recently introduced by Parker et al. (2013) was applied to identify the signatures of convergent evolution of *gpsA* versions after horizontal acquisition.

### Phylogenetic analysis of *Bartonella* species

In order to parsimoniously map HGT events to the evolutionary history of bartonellae, phylogenetic relationships among *Bartonella* species were inferred using standard phylogenetic and phylogenomic approaches as follows: *I. Phylogenomic analysis*: The proteomes of 23 annotated *Bartonella* genomes were downloaded, from which orthologous groups (OGs) were identified using OrthoMCL 2.0 (Li et al. 2003) with default parameters (BLASTP E-value cutoff = 1 × 10^−5^, percent match cutoff = 80%, MCL inflation parameter = 1.5). OGs that have exactly one member in each and every genome were isolated, resulting in 516 OGs. Members of each of these OGs were aligned in MAFFT (Katoh and Standley 2013) and refined in Gblocks 0.91b (Castresana 2000) to remove problematic regions. An optimal amino acid substitution model for each OG was computed in ProtTest (Darriba et al. 2011) using the BIC criterion. The 516 alignments were concatenated into one dataset, based on which a phylogenetic tree was reconstructed using the maximum likelihood (ML) method as implemented in RAxML (Stamatakis 2006) with 100 fast bootstrap replicates. *Bartonella tamiae* was used as an outgroup, with all other bartonellae treated as ingroup. *II. Standard phylogenetic analysis*: Five additional species could not be included in above approach, since their proteomes are not available from GenBank. To explore and confirm the phylogenetic positions of these *Bartonella* species, a separate analysis following previously outlined approaches (see above) was performed using six commonly used gene markers (Inoue et al. 2010; Mullins et al. 2013; Sato et al. 2012) from 28 *Bartonella* genomes (*B. tamiae* included in ingroup) and seven outgroup genomes which represent close sister genera of *Bartonella* (Table S1) (Gupta and Mok 2007; Guy et al. 2013).

### Experimental genotyping

In order to further explore the distribution of *Bartonella* clades with HGT derived metabolic genes in blood-feeding insects, we screened a global sampling of 21 species of Siphonaptera and Hippoboscoidea for *gpsA* sequences (Table 2). All of these samples had been positive for *Bartonella gltA* gene and 16S rRNA detection by PCR in previous analyses (Morse et al. 2012; Morse et al. 2013). Genomic DNA was extracted from each individual specimen, using the DNeasy Blood & Tissue Kit (Qiagen Sciences Inc., Germantown, MD, USA), following the animal tissue protocol. The quality and concentrations of DNA were assessed with a NanoDrop spectrometer (Thermo Fisher Scientific, Wilmington, DE, USA). Bacterial *gpsA* diversity was assessed by amplification of *gpsA* genes from each sample using specific primers and reaction conditions: *Helicobacter*-derived *gpsA* (He) (see Results) forward: 5′-ATG AAA ATA ACA RTT TTT GGW GGY GG-3′, reverse: 5′-TTA ATA CCT TCW GCY ACT TCG CC-3′; Enterobacteriales-derived *gpsA* (Ar/Se) forward: 5′-GGT TCT TAT GGY ACY GCW TTA GC-3′, reverse: 5′-TAR ATT TGY TCG GYA ATT GGC ATT TC-3′. Subsequent TA cloning (if applicable) was performed to isolate amplicons. Based on previous studies of the microbial diversity of bat flies, we expect a subset of species to harbor *Arsenophonus* species as an endosymbiont (Morse et al. 2013). In these species we specifically targeted *Arsenophonus gpsA* for comparative purposes. Sequence analysis and phylogenetic analysis followed the standard protocols described above.

## Results

The initial discovery screen revealed that several fundamental genes involved in the phospholipid pathway show patterns of repeated homologous replacements from identifiable sources outside Alphaproteobacteria, and/or gene loss (Figure 1). These genes are: 1) the *gpsA* gene, which is a minimal core gene encoding NAD(P)H-dependent glycerol-3-phosphate dehydrogenase, an enzyme that is essential to the synthesis of bacterial membrane lipids; 2) the *glpK* gene (glycerol kinase), which encodes an enzyme that is located in the cell membrane and catalyzes the Mg^2+^-ATP-dependent phosphorylation of glycerol to G3P (Glycerol-3-Phosphate); and 3) the Glp system (encoded by genes *glpS-T-P-Q-U-V*), an ABC transporter that is responsible for importing extracellular glycerol (Ding et al. 2012).

**Figure 1.**
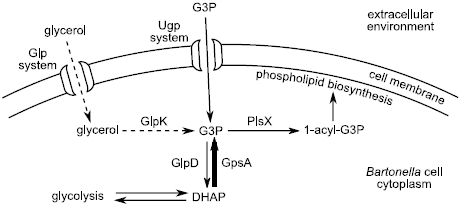
Role of GpsA in *Bartonella* phospholipid biosynthesis. Part of the alphaproteobacterial phospholipid biosynthesis pathway is illustrated based on (Cronan 2003; Pereto et al. 2004; Spoering et al. 2006; Yeh et al. 2008), and the KEGG pathway entry bhe00564 (glycerophospholipid metabolism in *B. henselae*). The illustration highlights the three possible paths of obtaining G3P (Glycerol-3-phosphate). The dashed lines represent the reactions affected by ancient gene losses; and the bold line represents the reaction affected by one ancient gene loss followed by three independent horizontal regains in the evolutionary history of *Bartonella*.

### Loss of *glpK* and Glp system precedes loss of *gpsA*

Results of BLAST-based and phylogenetic approaches reveal a pattern of additional gene losses in a core alphaproteobacterial metabolic pathway. Specifically, key genes involved in the glycerol pathway are ancestrally lost in the bartonellae. Only the *B. tamiae* genome, the most basal *Bartonella* species, contains *glpK*. However, phylogenetic analysis places this copy closely related to Enterobacteriaceae, which is suggestive of a horizontal origin. This, together with the complete absence of *glpK* and the Glp system in extant eubartonellae, supports a loss of the glycerol pathway at the base of all currently known bartonellae, preceding the *gpsA* loss (Figure 2). The absence of GlpK and the Glp system precludes the ability of eubartonellae to utilize extracellular glycerol as a source of G3P (Figure 1). No other functional homologs of these genes are known, or have been found in our genomic analysis.

**Figure 2.**
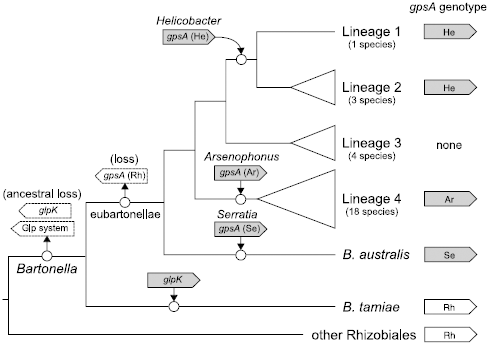
Losses and gains of *gpsA* and metabolically related genes in the evolutionary history of *Bartonella*. Schematically illustrated relationships of *Bartonella* lineages based on phylogenomic and phylogenetic analyses of the 28 *Bartonella* species (Figure S1). Topology is congruent with a recent phylogenomic study (Guy et al. 2013). Major monophyletic lineages (Table 1) were collapsed into triangles. Branch lengths are not drawn to scale. The presence and origin of *gpsA* is indicated to the right of corresponding lineages. Horizontally acquired genes are indicated by grey boxes, while vertically inherited genes are indicated by white boxes. Horizontal gene transfer events are represented by incoming arrows, with the putative donor groups (if identifiable) labeled. Gene loss events are represented by outgoing arrows and boxes with dashed outlines. Phylogenetic positions of losses and gains are indicated by circles.

### Multiple origins of *Bartonella gpsA* genes

BLAST-based and phylogenetic analyses reveal four origins of *gpsA* genes among *Bartonella* genomes (Figure 2, Figure S1). Known functional equivalents (not homologs) of bacterial GpsA (G3PDH), such as archaeal EgsA (G1PDH) are not present in any of the genomes (Koga et al. 1998). No *Bartonella* species has more than one copy of *gpsA*. Only the earliest diverging *B. tamiae* (Guy et al. 2013) contains a *gpsA* gene close to those of other Rhizobiales (Figure 3A). Specifically, *B. tamiae* is placed as a sister group to Brucellaceae (*Brucella* and *Ochrobactrum*), within the Rhizobiaceae (*Rhizobium, Agrobacterium* and *Sinorhizobium*). This topology mirrors our current knowledge of Rhizobiales and *Bartonella* evolution (Gupta and Mok 2007; Munoz et al. 2011).

**Figure 3.**
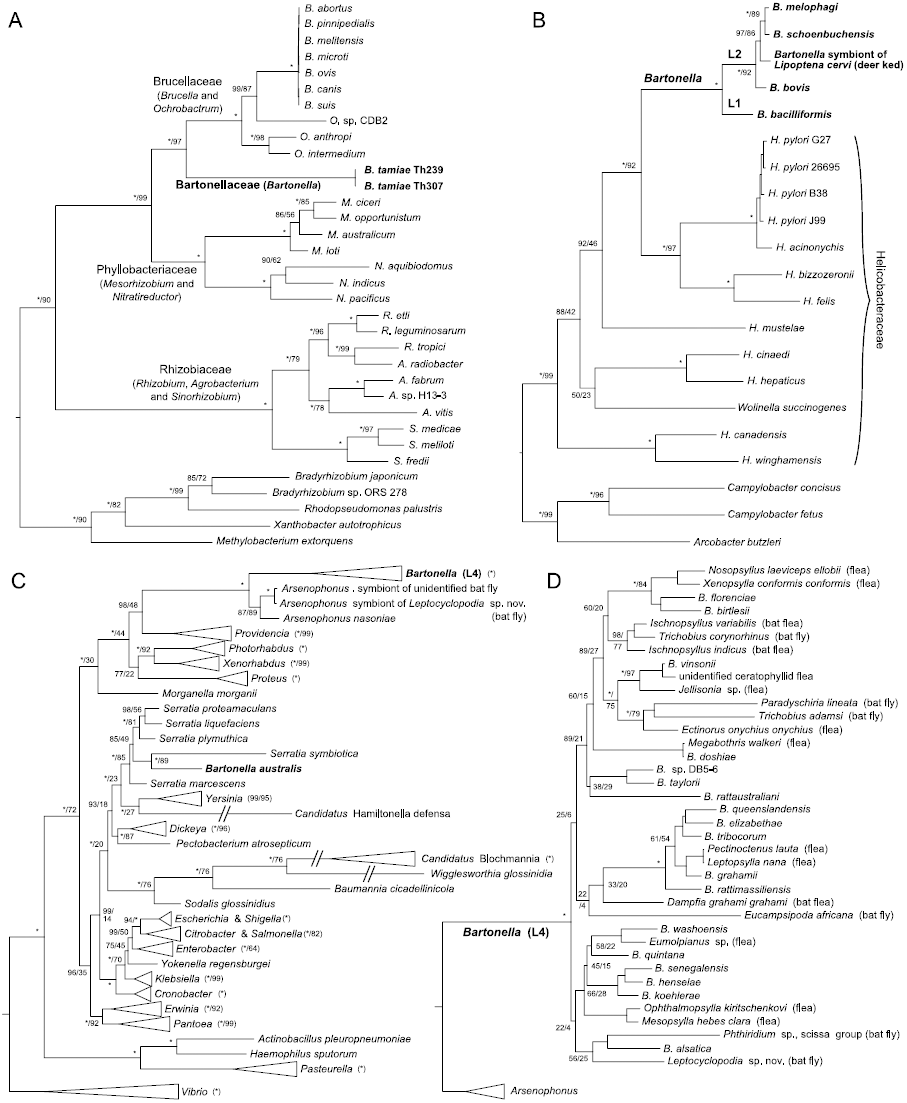
Phylogenies of different versions of *gpsA*. Trees were reconstructed using Bayesian inference as implemented in MrBayes. Node labels (x/y) represent Bayesian posterior probabilities (x %) computed in MrBayes and maximum-likelihood bootstrap support values (y % out of 1000 replicates) computed in RAxML. Asterisks (*) indicate 100% support. *Bartonella* clades are denoted in bold font. (**A**) Vertical inheritance history of *gpsA* (Rh) in Rhizobiales. Families Bartonellaceae, Brucellaceae, Phyllobacteriaceae and Rhizobiaceae are placed as ingroups and the other Rhizobiales organisms as outgroups, according to Gupta and Mok (2007). (**B**) Horizontal transfer of *gpsA* (He) from *Helicobacter* to L1 and L2 *Bartonella* (including an experimentally verified deer ked sample). The tree is rooted at the common ancestor of Helicobacteraceae and Campylobacteraceae, according to Gupta (2006). (**C**) Horizontal transfers of *gpsA* (Ar) from *Arsenophonus* to L4 *Bartonella*, and that of *gpsA* (Se) from *Serratia* to *B. australis*. The tree includes recipient *Bartonella* species, representative Enterobacteriales groups and two *Arsenophonus*-positive bat fly samples sequenced in this study. It is rooted to Vibrionales according to Williams et al. (2010). Monophyletic groups are collapsed in triangles with nodal support values labeled to the right. Long branches are truncated and indicated by two slashes (//). (**D**) Post-transfer evolutionary history of *Arsenophonus*-derived *gpsA* (Ar) in L4 *Bartonella*. This is an expansion of the L4 *Bartonella* clade in C. Experimentally verified insect samples are indicated by the insect names. Nodal support values of derived clades are omitted.

*Bartonella bacilliformis* (lineage 1), and all members of lineage 2 (*B. bovis, B. melophagi* and *B. schoenbuchensis*) possess *a gpsA* that nests strongly supported within a genus of Epsilonproteobacteria, namely *Helicobacter* (Figure 3B). In the phylogenetic analysis all *Helicobacter*-derived *gpsA* genes [*gpsA* (He)] form a strict monophyletic group, with lineage 1 and lineage 2 split at the base. Its immediate sister group is a clade of four *Helicobacter* species (Figure 3B). The general structure of the Helicobacteraceae clade resembles the species tree of this family from previous studies (Dewhirst et al. 2005; Gupta 2006; Munoz et al. 2011).

Lineage 3 bartonellae (*B. clarridgeiae, B. rochalimae, B. sp.* 1-1C and *B. sp.* AR 15-3) lack any identifiable homolog of *gpsA*.

*B. australis* and all members of lineage 4 (Table 1) possess *gpsA* genes which were captured from the gammaproteobacterial Enterobacteriales. These *gpsA* genes were transferred in separate instances, as *B. australis gpsA* has high sequence similarity and phylogenetic affiliation with *Serratia* species [*gpsA* (Se)], whereas all available representatives of lineage 4 contain an *Arsenophonus*-derived *gpsA* gene [*gpsA* (Ar)] (Figure 3C, D). Specifically, analyses reveal that all lineage 4 *Bartonella gpsA* genes form a monophyletic group that is sister to extant *Arsenophonus* species. Together they are nested within a clade including *Arsenophonus’* closest sister groups: *Providencia, Photorhabdus, Xenorhabdus*, and *Proteus*. On the other hand, *B. australis gpsA* nests within the *Serratia* clade, with its closest sister group being *Serratia symbiotica*.

### Genomic environments of the *gpsA* genes support one ancestral loss and three individual transfers at the base of major *Bartonella* lineages

#### Rhizobiales-derived gpsA (Rh)

The *gpsA* (Rh) gene (Figure 5A) is located within a gene cassette that typically contains five tandemly arranged genes in the genome of *Bartonella tamiae* and those of close sister groups of *Bartonella*, including *Brucella, Ochrobactrum, Mesorhizobium, Agrobacterium, Rhizobium* and *Sinorhizobium*. In all other *Bartonella* genomes, this cassette is still present, but consistently rhizobial *gpsA* and its immediate downstream open reading frame (ORF) (Ycil-like protein CDS) are absent in all eubartonellae. Instead, this space is occupied by sequences without identifiable ORFs. No sequence similarity can be detected between those sequences and the original contents. The above-described pattern is consistent with a single ancestral loss of *gpsA* followed by three gains (see below, Figure 2), each of which coincides with the current lineage classification of bartonellae (Engel et al. 2011). From all known bartonellae, lineage 3 is the only clade in which all species are not only missing the *glpK* and the *glp* genes, but it also never re-gained *gpsA* (Figure 2).

#### Helicobacter-derived gpsA (He)

The original *gpsA* (He) (Figure 5B) is residing in a genomic environment that is highly variable among *Helicobacter* species. In most cases, it is upstream of the *glyQ* (glycyl-tRNA synthetase subunit alpha) gene. In lineage 1 and 2 *Bartonella* genomes, only the *gpsA* gene seems to have been transferred (Figure 2), without its upstream and downstream neighbors from *Helicobacter*. The gene is located in a genomic locus, where the upstream side is a group of four ORFs ending with the *hisS* (histidyl-tRNA synthetase) gene. The downstream side of *gpsA* in lineage 2 *Bartonella* genomes is an rRNA operon, which is typically present in all *Bartonella* genomes as two to three copies (Guy et al. 2013; Viezens and Arvand 2008). The horizontal transfer of *gpsA* (He) into lineage 1 and 2 seems to have interrupted an ORF present in all bartonellae, which in lineage 2 bartonellae is still present with a residual sequence (Figure 5B). Phylogenetic analysis of this ORF sampled across all bartonellae mirrors current hypotheses of *Bartonella* species evolution (Guy et al. 2013).

#### Arsenophonus-derived gpsA (Ar)

In the genomes of *Arsenophonus* and other Enterobacteriaceae, the original *gpsA* (Ar) gene (Figure 5C) resides within a cassette of four genes (*secB-gpsA-cysE-cspR*) right downstream of the O-antigen gene cluster, a frequently horizontally transferred structure (Ovchinnikova et al. 2013; Wildschutte et al. 2010). In lineage 4 *Bartonella* genomes, *gpsA* (Ar) seems to have been transferred singularly into a genomic region that is present in all bartonellae. Upstream of it is a *cyo* operon (*cyoA, B, C, D*) encoding the cytochrome *o* ubiquinol oxidase, a component of the aerobic respiratory chain (Reva et al. 2006). Downstream is a cluster of three genes (*fabI1-fabA-fabB*) that are essential in fatty acid biosynthesis (Campbell and Cronan 2001).

#### Serratia-derived gpsA (Se)

In *Serratia, gpsA* is located in the homologous gene cassette to other Enterobacteriaceae. In the *B. australis* genome, *gpsA* was cotransferred with two other genes of the donor cassette (*grxC-secB-gpsA*), and resides in a region that is highly variable among *Bartonella* species. However, the structures upstream and downstream of this highly variable region are relatively constant in all *Bartonella* genomes (Figure 5D). Notably, several house-keeping genes involved in lipid metabolism (*plsX, fabH, accB, accC*, and *glpD*) (Campbell and Cronan 2001; Cronan 2003) are located proximal to this region.

### Other genes in the phospholipid pathway

Meanwhile, an alternative route of G3P acquisition is intact: The Ugp system (encoded by the operon *ugpB-A-E-C*), an ABC transporter that imports G3P into cells (Brzoska et al. 1994), is present in all extant *Bartonella* species. Phylogenetic analysis reveals that this operon is vertically transmitted in the eubartonella (Figure S3).

G3P’s utilization in the phospholipid biosynthesis pathway is mediated by PlsX (G3P acyltransferase), which converts G3P into 1-acyl-G3P for subsequent steps. PlsX-deficient *E. coli* strains cannot synthesize a cell membrane (Bell 1974). All *Bartonella* species maintain one copy of this gene. The phylogenetic tree of *plsX* mirrors the species tree of *Bartonella* (Figure S4).

### Molecular evolution analyses

All genes tested in this analysis maintain an ORF (regardless of horizontal or vertical origin). Analyses testing for selective pressure along branches and among sites were carried out on trees of individual *gpsA* gene families and other metabolically related genes (Table S2). The following general patterns that apply to all three *gpsA* families were observed: There are clear signatures of global stabilizing selection operating on *gpsA* genes, including on branches leading up to the nodes representing the transfers from putative donor groups (*Helicobacter, Arsenophonus* and *Serratia*) to stem-*Bartonella* of the major lineages (L1+L2, L4, and *B. australis*, respectively). Strong stabilizing selection was also observed in horizontally acquired and subsequently vertically transmitted copies of *Bartonella gpsA*. Within the *Bartonella* clades (L1+L2, L4 and *B. australis* alone) that represent the evolutionary history of *gpsA* after acquisition, the ω values are significantly elevated compared to the tree backgrounds (Table S2), suggesting an accelerated rate of evolution after horizontal acquisition.

Representative protein sequences of each version of *gpsA* were aligned to the *Coxiella burnetii* sequence, whose tertiary structure has been experimentally verified (Figure 4). The sequence similarity among versions is generally low. However, functional motifs and their adjacent sites exhibit strong conservation across all *gpsA* sequences. None of these sites were predicted to be under positive selection. The majority of sites that were detected to be under significant positive (diversifying) selection [(27/31), prob >95%] have been identified in the *gpsA* of lineage 4 *Bartonella* (Table S2, Figure 4).

**Figure 4.**
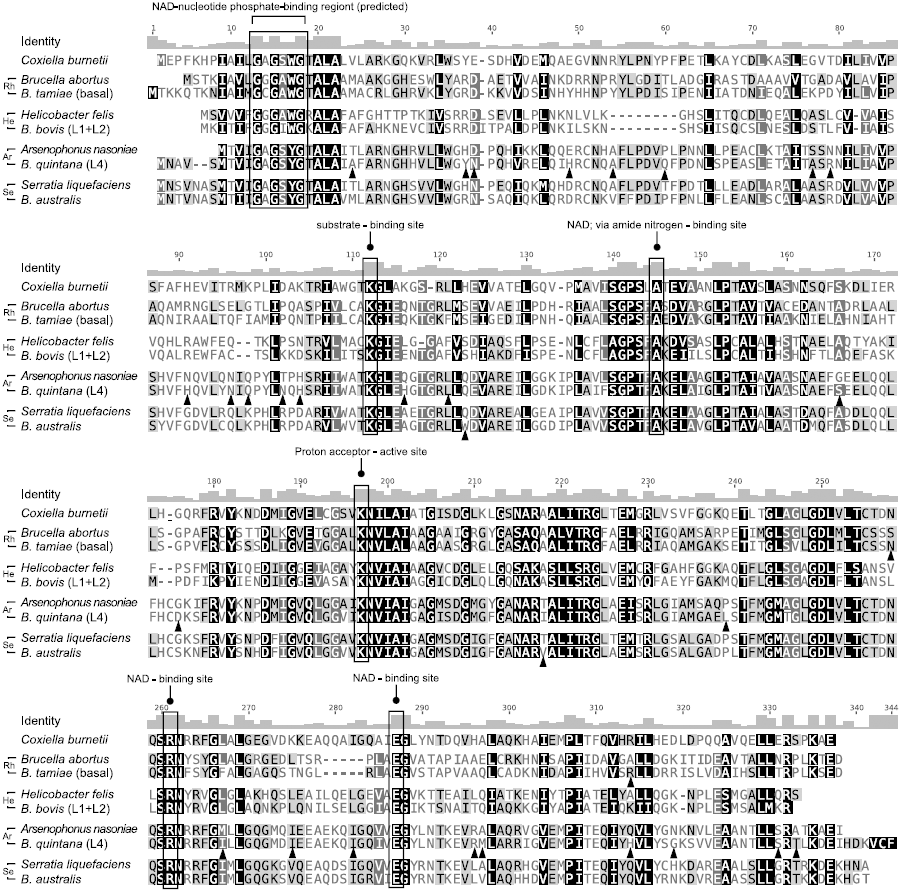
Comparison of protein sequences of different GpsA versions. Alignment of full-length GpsA protein sequences to *Coxiella burnettii*. Nucleotide positions are shaded by similarity from low (light) to high (dark) on a grayscale. GpsA proteins are aligned in pairs with a representative sequence from the donor group and its *Bartonella* counterpart. Functional sites and motifs are boxed, as recorded in UniProtKB. Significantly positively selected sites predicted by BEB are indicated by solid triangles.

**Figure 5.**
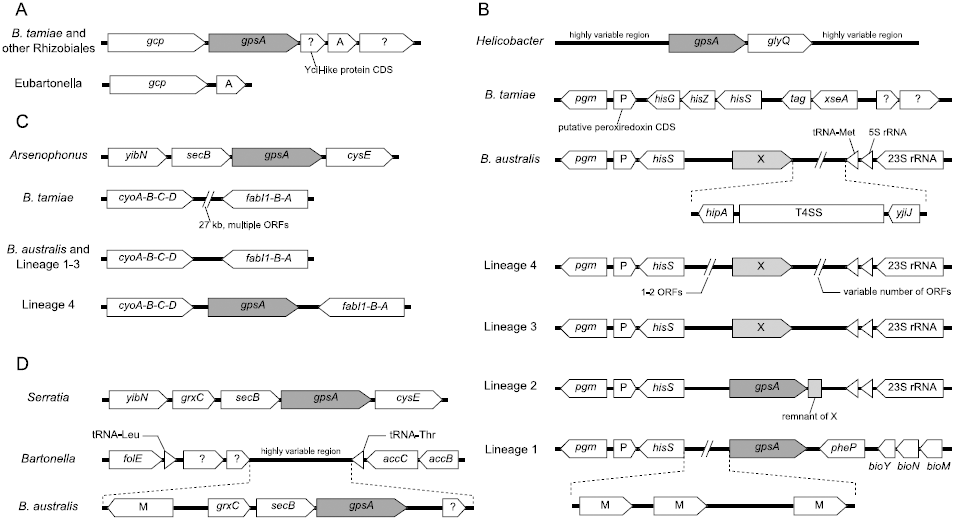
Genomic contexts of different versions of *gpsA* in *Bartonella* and other bacterial groups. Genes are represented by boxes. Tandemly arranged genes that share the same symbol (a putative operon) are merged. Lengths of genes and intergenic regions are not drawn to scale. Open reading frames (ORFs) annotated as hypothetical genes are either indicated by “?”, or by single letters (e.g., “M” and “X”, see below). Scenarios illustrated in the panels are: (**A**) Loss of *gpsA* (Rh) in stem eubartonella. (**B**) HGT of *gpsA* (He) from *Helicobacter* to the common ancestor of L1 and L2 *Bartonella*. (**C**) HGT of *gpsA* (Ar) from *Arsenophonus* to stem L4 *Bartonella*. (**D**) HGT of *gpsA* (Se) from *Serratia* to *Bartonella australis*. “X” represents an ORF that is disrupted by the insertion of *gpsA* (He). “M” represents a multi-copy ORF that exists only in *B. bacilliformis* and *B. australis* genomes.

Applying the statistical methods outlined by Parker et al. (2013) resulted in no significant signature of convergent evolution in horizontally transferred *gpsA* versions across *Bartonella*.

All other genes in the phospholipid pathway are under strong stabilizing selection across all lineages in the bartonellae (Table S2).

### Phylogenetic analyses of *Bartonella*

Phylogenomic analysis of *Bartonella* core genomes (23 species) and of selected genes (28 species) recovers previously identified clades, and relationships. Topologies of ingroup bartonellae (eubartonellae) mirror results from the recent analysis of Guy et al. (2013) (Table 1, Figure S1). Although there is some controversy about the relationship of L1 and L2 bartonellae to each other (Engel et al. 2011; Guy et al. 2013), the observed pattern of ORF disruption by the *Helicobacter*-derived *gpsA* (see above) provides a solid piece of evidence of a single transfer event, and a shared derived ancestry of L1 and L2. *Bartonella tamiae* occupies a strongly supported ancestral position to all other bartonellae in the standard phylogenetic analysis. We recovered a basal position for *Bartonella australis* in eubartonellae in both analyses, and the evolutionary sequence of lineage-specific diversification is strongly supported on all nodes.

### Experimental genotyping of *Bartonella gpsA*

*GpsA* sequences were successfully recovered from most of our samples of blood-feeding insects (Table 2). One bat fly sample contained two *gpsA* copies – one copy of the known, and expected *Arsenophonus* endosymbiont of this group (Morse et al. 2013), and one copy of *Bartonella gpsA* (Ar). *Helicobacter*-type *gpsA* [*gpsA* (He)] was detected in a deer ked (*Lipoptena cervi*) sample, and is phylogenetically nested within the L2 *Bartonella* clade (Figure 3B). *Arsenophonus-type gpsA* [*gpsA* (Ar)] was found in most fleas (Siphonaptera) and in all bat flies (Nycteribiidae and Streblidae), increasing the number of known bartonellae vector species for both fleas, and bat flies. All of these are distributed within the L4 *Bartonella* clade, which shows species groupings generally consistent with previous analyses (Figure S1). Flea and bat fly host affiliations scatter throughout previously known subclades. No *Serratia*-type *gpsA* [*gpsA* (Se)] was detected in any of our samples (*Bartonella australis*). The currently known distribution of *B. australis* is restricted to Australia (Fournier et al. 2007), and doesn’t overlap with our sampling. Several fleas did not yield any *gpsA* gene, despite being positive for *Bartonella gltA* and 16S rRNA pertaining to lineage 3.

**Table 2.**
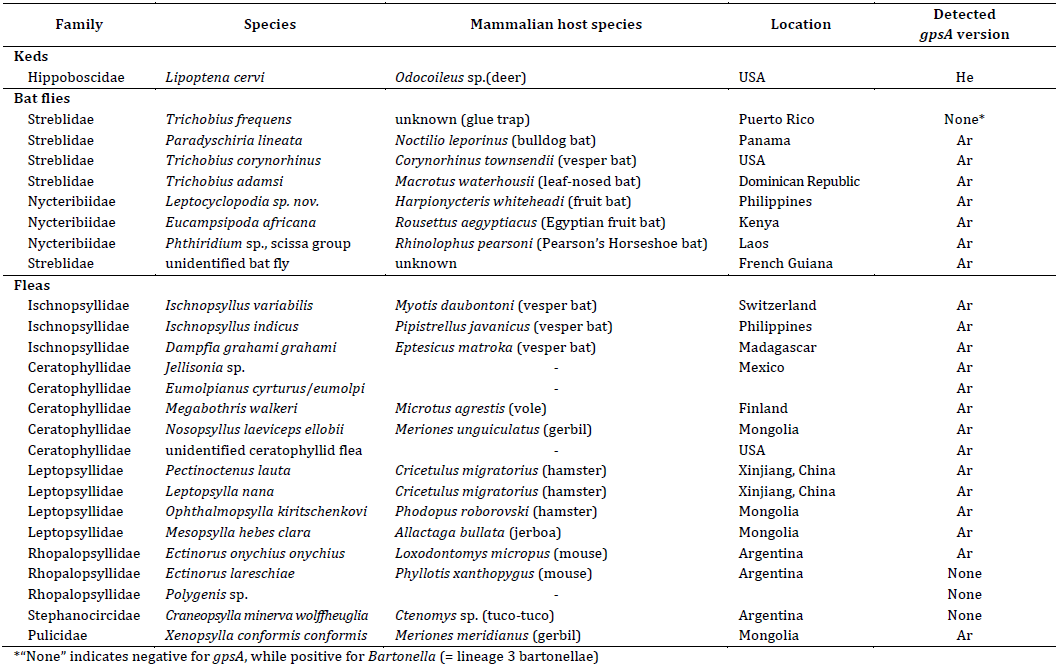
PCR-verified *gpsA* types in *Bartonella*-positive insect samples

| Family | Species | Mammalian host species | Location | Detected *gpsA* version |
| --- | --- | --- | --- | --- |
| **Keds** |  |  |  |  |
| Hippoboscidae | *Lipoptena cervi* | *Odocoileus* sp.(deer) | USA | He |
| **Bat flies** |  |  |  |  |
| Streblidae | *Trichobius frequens* | unknown (glue trap) | Puerto Rico | None* |
| Streblidae | *Paradyschiria lineata* | *Noctilio leporinus* (bulldog bat) | Panama | Ar |
| Streblidae | *Trichobius corynorhinus* | *Corynorhinus townsendii* (vesper bat) | USA | Ar |
| Streblidae | *Trichobius adamsi* | *Macrotus waterhousii* (leaf-nosed bat) | Dominican Republic | Ar |
| Nycteribiidae | *Leptocyclopodia* sp. nov. | *Harpionycteris whiteheadi* (fruit bat) | Philippines | Ar |
| Nycteribiidae | *Eucampsipoda africana* | *Rousettus aegyptiacus* (Egyptian fruit bat) | Kenya | Ar |
| Nycteribiidae | *Phthiridium* sp., scissa group | *Rhinolophus pearsoni* (Pearson's Horseshoe bat) | Laos | Ar |
| Streblidae | unidentified bat fly | unknown | French Guiana | Ar |
| **Fleas** |  |  |  |  |
| Ischnopsyllidae | *Ischnopsyllus variabilis* | *Myotis daubontoni* (vesper bat) | Switzerland | Ar |
| Ischnopsyllidae | *Ischnopsyllus indicus* | *Pipistrellus javanicus* (vesper bat) | Philippines | Ar |
| Ischnopsyllidae | *Dampfia grahami grahami* | *Eptesicus matroka* (vesper bat) | Madagascar | Ar |
| Ceratophyllidae | *Jellisonia* sp. | - | Mexico | Ar |
| Ceratophyllidae | *Eumolpianus cyrturus/eumolpi* | - |  | Ar |
| Ceratophyllidae | *Megabothris walkeri* | *Microtus agrestis* (vole) | Finland | Ar |
| Ceratophyllidae | *Nosopsyllus laeviceps ellobii* | *Meriones unguiculatus* (gerbil) | Mongolia | Ar |
| Ceratophyllidae | unidentified ceratophyllid flea | - | USA | Ar |
| Leptopsyllidae | *Pectinotenus lauta* | *Cricetulus migratorius* (hamster) | Xinjiang, China | Ar |
| Leptopsyllidae | *Leptopsylla nana* | *Cricetulus migratorius* (hamster) | Xinjiang, China | Ar |
| Leptopsyllidae | *Ophthalmopsylla kiritschenkovi* | *Phodopus roborovski* (hamster) | Mongolia | Ar |
| Leptopsyllidae | *Mesopsylla hebes clara* | *Allactaga bullata* (jerboa) | Mongolia | Ar |
| Rhopalopsyllidae | *Ectinorus onychius onychius* | *Loxodontomys micropus* (mouse) | Argentina | Ar |
| Rhopalopsyllidae | *Ectinorus lareschiae* | *Phyllotis xanthopygus* (mouse) | Argentina | None |
| Rhopalopsyllidae | *Polygenis* sp. | - |  | None |
| Stephanocircidae | *Craneopsylla minerva wolffheuglia* | *Ctenomys* sp. (tuco-tuco) | Argentina | None |
| Pulicidae | *Xenopsylla conformis conformis* | *Meriones meridianus* (gerbil) | Mongolia | Ar |

\*"None" indicates negative for *gpsA*, while positive for *Bartonella* (= lineage 3 bartonellae)

## Discussion

### Ancestral intracellularity in *Bartonella*

Given the conserved nature of the bacterial phospholipid pathway, the ancestral loss of *glpK* and the Glp system in stem bartonellae, followed by a *gpsA* loss at the base of eubartonellae likely created an ancestral population of bartonellae unable to use either glycerol, or glucose metabolites (Figure 1). Our analyses therefore suggest that the ancestors of extant eubartonellae relied directly on G3P, which can be imported into the cytoplasm by the Ugp system that remains intact in all bartonellae. G3P is known to be an intermediate metabolite of a strictly intracellular biochemical pathway in prokaryotic and eukaryotic cells, and doesn’t occur stably in the extracellular environment (e.g., blood) (Cronan 2003; Spoering et al. 2006). Therefore, G3P capture and utilization by ancestral bartonellae were likely accomplished by cytoplasmic associations to a living cell. Previous research has shown that in the absence of readily available G3P, the ability of *gpsA* mutant bacteria to form cell membranes is severely compromised, resulting in the cessation of cell growth. (Bell 1974; Cronan and Bell 1974). This functional peculiarity may explain the slow growth of the gpsA-less lineage 3 bartonellae on blood agar, and their more successful isolation in living cell lines (Heller et al. 1997; Podsiadly et al. 2007). Specifically, the four extant representatives of lineage 3 bartonellae (*B. rochalimae, B. clarridgeiae, B.* sp. 1-1C, and *B.* sp. AR 15-3) are possibly the surviving representatives of an ancient lineage, as they are still lacking *glpK*, the Glp system, as well as *gpsA*. The ubiquitous loss of these important genes in the phospholipid pathway prior to the evolution of extant eubartonellae certainly suggests that bartonellae had an early intracellular beginning.

### Functional importance of the acquired *gpsA* genes

Our results provide strong evidence that bartonellae *gpsA* was acquired from three independent prokaryotic sources outside of alpha-proteobacteria after a single initial loss at the base of the eubartonellae lineage. The repeated retentions of HGT-derived *gpsA* in the *Bartonella* genomes confirm the functional importance of *gpsA*, in the context of the loss-and-regain hypothesis of Doolittle et al. (2003). Based on an array of studies related to HGT, it has been hypothesized that genes that are selectively advantageous in the new organisms have a higher probability of being retained (Kuo and Ochman 2009). Results suggest vertical inheritance and global stabilizing selection after *gpsA* transfer in *Bartonella* lineages, as well as the maintenance of open reading frames in all transferred genes. Taken together with the previously confirmed expression of *gpsA* in bartonellae (Omasits et al. 2013; Saenz et al. 2007) the above evidence supports the functionality of the *gpsA* genes after transfer. Furthermore, the protein sequence alignment shows that all functionally important sites are conserved among the HGT derived *gpsA* versions (Figure 3). This, combined with the overall stabilizing selection on horizontally transferred *gpsA* genes (see Results) imply that the acquired genes are likely to have inherited their original functional role in the biosynthesis of bacterial membrane lipids. However, the significant elevation of evolutionary rates in all three acquired genes and the detection of specific sites under positive selection suggest that amid the overall stabilizing selection, the genes may still undergo functional evolution to adapt to bartonellae-specific metabolic pathways. Moreover, although different *gpsA* genes were inserted into distinct genomic loci, it is notable that their genomic contexts typically include clusters of genes that are involved in bacterial lipid metabolism (see Results; Figure 5). This suggests that they have been integrated into the existing transcriptional regulation machinery of lipid metabolic genes, which may have facilitated their retention (Lercher and Pal 2008). Bartonellae *gpsA* mutants are known to result in an abacteremic phenotype (Saenz et al. 2007), pointing to the importance of functional *gpsA* in pathogenicity and hematogenous spread. Therefore it is possible that the horizontal re-acquisitions of functional *gpsA* facilitated the hematogenous spread of bartonellae to diverse hosts through blood-feeding vectors, as confirmed for the majority of extant bartonellae.

### Ancestral host-associations of bartonellae

For two prokaryotic organisms to exchange genetic material requires direct (e.g., conjugation, transformation) or very proximal contact (e.g., phages) in an appropriate environment (Frost et al. 2005; Polz et al. 2013). Therefore, the identified cases of HGT provide clues to infer past shared ecological connections between *Bartonella* and other bacteria as well as their ancestral putative hosts.

In line with this argumentation, we suggest that the two transfers from Gammaproteobacteria are more likely to have occurred in an arthropod. Specifically, the *Arsenophonus*-derived transfer likely stems from endosymbionts exclusively associated with arthropods. *Arsenophonus*, the immediate well supported sister group of the bartonellae *gpsA*, is ecologically versatile, and widely distributed among arthropods, including blood-feeders, such as ticks (Ixodidae), keds (Hippoboscidae) and bat flies (Nycteribiidae) (Morse et al. 2013; Novakova et al. 2009; Trowbridge et al. 2006). Among blood-feeding parasites, *Arsenophonus* species are still primary endosymbionts of extant hippoboscoid flies, which are among the confirmed insect vectors of bartonellae of lineages 4 and 2 (Halos et al. 2004; Morse et al. 2013). Therefore, we propose that hippoboscoid flies may have already been among the ancestral blood-feeding arthropod hosts of bartonellae providing a biological reservoir conducive for horizontal transfer of genes. Interestingly, the gamma-proteobacterial *Arsenophonus* has never been detected in extant Siphonaptera, the insect order with the most common and speciose *Bartonella* vectors (Chomel et al. 2009; Pulliainen and Dehio 2012; Tsai et al. 2011). Yet all lineage 4 bartonellae transmitted by fleas carry the *Arsenophonus*-derived *gpsA* (Ar) (Figure 3C). These facts imply that the *Arsenophonus*-derived transfer of *gpsA* (Ar) to stem L4 *Bartonella* likely did not occur in fleas, but that use of Siphonaptera are likely evolutionarily derived vectors of lineage 4 bartonellae.

The *gpsA* transfer from the enterobacterial *Serratia* involves *Bartonella australis*, which at present seems to be the only representative of its lineage, although this may change with better sampling coverage. *Serratia* species colonize diverse habitats, including plants, insects and vertebrates (Grimont and Grimont 2006). Given this wide, and largely underexplored host range, it is difficult to pinpoint a specific ancestral host for the HGT exchange of the *Serratia*-derived *gpsA* in *Bartonella*. However, the *Serratia*-derived *gpsA* of *B. australis* strongly nests within a monophyletic clade containing *Serratia symbiotica*, a known endosymbiont of aphids (Aphidoidea) (Figure 3C). In insects *Serratia* may function as pathogen or symbiont, and have been shown to invade the hemocoel and the intestinal tract (Grimont and Grimont 2006). In mammals, infection often is opportunistic, and rarely systemic (unless previous immunosuppression exists) (Grimont and Grimont 2006; Mahlen 2011). Therefore, it is conceivable that an insect host was involved in this transfer too.

In contrast, we suggest that the horizontal integration of *Helicobacter*-derived *gpsA* into a *Bartonella* genome likely took place in a mammalian host, especially given that bartonellae *gpsA* (He) is firmly nested within the *Helicobacter* clade. *Helicobacter* are predominantly mammalian pathogens (Rogers 2012; Whary and Fox 2004), whose hosts overlap well with the known host range of *Bartonella* species. In their hosts helicobacter-type bacteria typically colonize the gastrointestinal tract and liver, causing peptic ulcers, chronic gastritis, cancer and other diseases. Meanwhile, blood is a secondary site for some species, where they adhere to erythrocytes, which is also the dominant infection site of bartonellae (Dubois and Boren 2007; Whary and Fox 2004).

It is important to note that the timing of each inferred horizontal transfer coincides with the subsequent speciation of extant bartonellae lineages, yet it is difficult to assess exactly when these events happened on an evolutionary scale. However, the transferred *gpsA* genes exclusively affect eubartonellae (= every lineage after *B. tamiae*), and the transferred genes are still closely related to extant prokaryotic groups, allowing donor identification. This could be in part due to strong functional constraints, but may also point to an evolutionarily recent HGT relative to the total lineage age. This would support a picture of a more recent diversification with invertebrates and mammals, as suggested by the hypothesis of “explosive radiation” of bartonellae by other authors (Engel et al. 2011; Guy et al. 2013).

### Extant host associations

Experimental *Bartonella* genotyping and subsequent phylogenetic analyses recover expected lineages given currently known host and vector ranges (Halos et al. 2004; Morse et al. 2012) (Table 2), and confirm predictions of *gpsA* origin based on phylogenomic analyses. Specifically, bartonellae of deer keds (*Lipoptena cervi*, Hippoboscidae), which are known vectors of lineage 2 bartonellae contain *Helicobacter*-derived *gpsA* (He) (Figure 3B). A diverse sampling of flea and bat fly species shows *Arsenophonus*-derived *gpsA* (Ar), as is expected for vectors of L4 *Bartonella* species (Figure 3D). Some samples (e.g., *Megabothris walkeri*) closely cluster with known L4 *Bartonella* species (e.g., *B. doshiae*). Others, such as bartonellae from *Trichobius* species, appear to be phylogenetically distant from any established *Bartonella* subclades, suggesting putative novel species in bat flies, which has been proposed previously (Billeter et al. 2012; Morse et al. 2012). These findings call for more in-depth studies to characterize extant *Bartonella* diversity.

The overall topology of the L4 *gpsA* (Ar) tree (Figure 3D) does not mirror the phylogeny of either fleas (Whiting et al. 2008) or bat flies (Dittmar et al. 2006; Petersen et al. 2007), suggesting the absence of *Bartonella*-insect coevolution within these two groups. For bat flies, this is in contrast to their primary endosymbionts, which exhibit notable revolutionary patterns with their fly hosts (Hosokawa et al. 2012; Morse et al. 2013). Flea and bat fly bartonellae are interspersed among each other in the tree, implying frequent horizontal transmission of L4 bartonellae between insect vectors, and low host specificity.

More detailed coevolutionary analyses of mammal-*Bartonella* and insect-*Bartonella* relationships are warranted given these findings.

### Conclusions

Our study shows that the phospholipid pathway in *Bartonella* has been affected by gene losses and gains throughout their evolution. Specifically, *glpK*, the Glp system, and *gpsA* were lost, but only *gpsA* genes were reacquired in eubartonellae by three independent horizontal transfers from Gamma-, and Epsilonproteobacteria. Results from this discovery-based study indicate a key impact of HGT on the ecological and functional evolution in bartonellae.

## Acknowledgements

This work was supported by the National Science Foundation [DEB 1050793 to K.D.].

## Supplementary Information

Supplementary Figures S1-S4, Tables S1-S2

